# HERO: A hierarchy-aware analysis pipeline for reducing and refining whole-brain atlas-mapped cellular datasets

**DOI:** 10.64898/2026.07.02.736093

**Authors:** Ava L. Shipman, Samuel W. Centanni

**Affiliations:** Department of Translational Neuroscience, Wake Forest University School of Medicine, Winston-Salem, NC, USA

**Author notes:** **Correspondence:** Samuel W. Centanni, Ph.D., Wake Forest University School of Medicine, Department of Translational Neuroscience, Medical Center Boulevard, Winston-Salem, NC 27157-1083.

**Keywords:** atlas, cell detection and counting, data reduction, data refinement, hierarchy analysis, light-sheet fluorescence microscopy, whole-brain

## Abstract

Advances in high-throughput mesoscale microscopy and machine learning-based image analysis pipelines have made unbiased whole-brain imaging widely accessible. However, translating the resulting atlas-mapped datasets into biologically meaningful results remains a substantial barrier owing to their sheer magnitude and complex hierarchical organization. Consequently, reporting structure and analysis methods vary widely across studies, undermining rigor and reproducibility. To address this, we developed a user-friendly data reduction workflow, HERO (Hierarchy-aware Expression Region Organization), designed to perform hierarchy-aware selection, refinement, ranking, and visualization of whole-brain cell detection datasets. The workflow is customizable to specific needs, requires minimal coding experience, and outputs transparent, curated results. HERO functions as a seamless plug-in within larger-scale whole-brain cell detection analysis pipelines, providing efficient, exploratory region selection to streamline subsequent statistical analyses and comparative evaluations. We applied HERO to whole-brain mapping datasets of afferents to the bed nucleus of the stria terminalis (BST) and the insular cortex (insula), areas we and others have previously characterized, giving a known connectivity baseline for evaluating the workflow. HERO provided top projecting regions consistent with prior tracing studies for the BST (e.g., amygdalar and hypothalamic regions) and the insula (e.g., somatosensory, autonomic, and interoceptive regions). Furthermore, the refinement step of HERO provides enriched regions originally obscured in the primary ranked outputs, such as memory- and emotion-associated structures. Beyond data reduction, HERO serves as an accessible tool for exploring and understanding complex whole-brain cell-detection datasets, particularly for researchers new to whole-brain imaging. This supervised data reduction process allows users to track analytical decisions, explore anatomical relationships, and gain deeper insight into their datasets beyond a single ranked list. In sum, HERO offers a standardized analysis workflow to reduce whole-brain cell detection datasets, transforming raw regional cell counts into curated results, advancing the effectiveness, interpretability, and accessibility of whole-brain imaging.

## 1 Introduction

Rodent whole-brain delipidation, imaging, atlas alignment, and cell-detection workflows have been transformative to neuroscience research and are now becoming more widely accessible and streamlined (Table 1) (Puchades et al., 2025, Wang et al., 2026). Generating high-quality 3D reconstructed data with mesoscale imaging creates opportunities to advance biomedical research by enabling integrated observation of dynamic biological processes. However, increasing accessibility has created a new bottleneck in data reduction and interpretation, leaving a fundamental gap in standardized, reproducible preprocessing methods for whole-brain cell detection data.

**Table 1.**
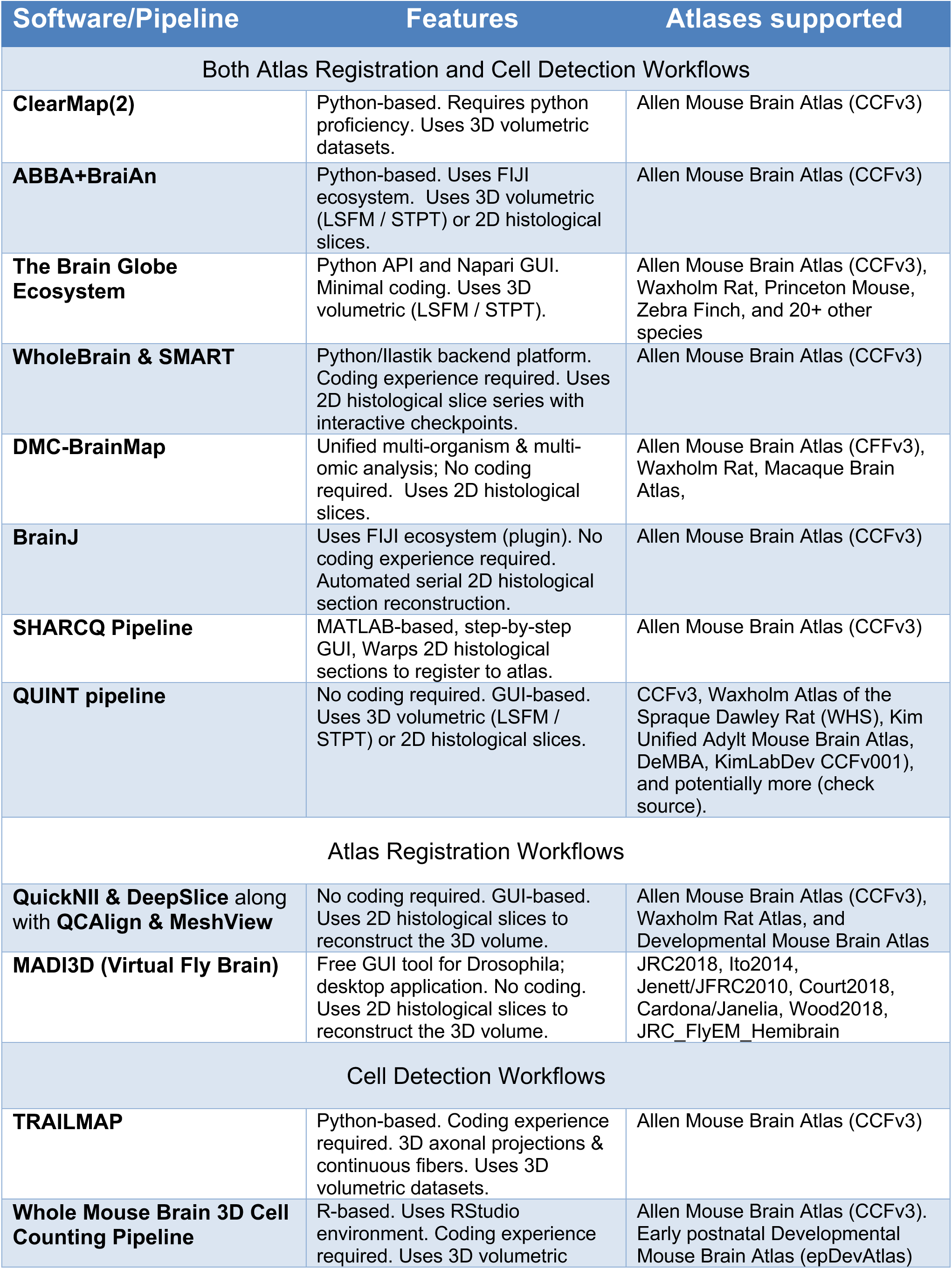
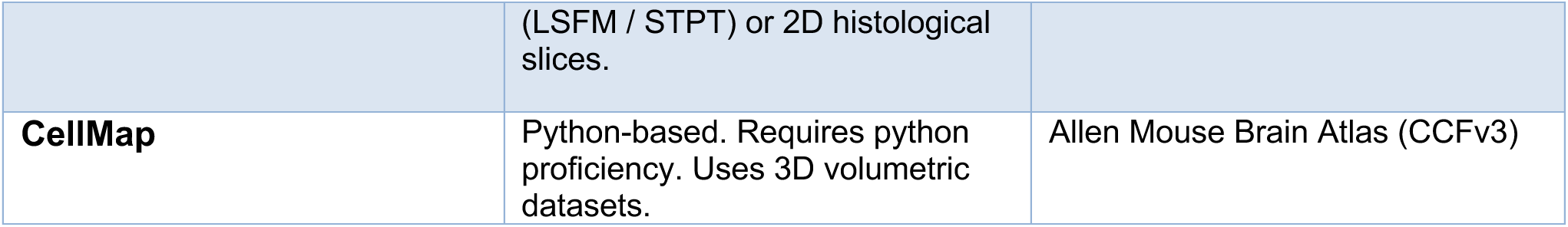
Examples of open-source whole-brain atlas registration and cell detection workflows.

| **Software/Pipeline** | **Features** | **Atlases supported** |
|---|---|---|
| Both Atlas Registration and Cell Detection Workflows |  |  |
| **ClearMap(2)** | Python-based. Requires python proficiency. Uses 3D volumetric datasets. | Allen Mouse Brain Atlas (CCFv3) |
| **ABBA+BraiAn** | Python-based. Uses FIJI ecosystem. Uses 3D volumetric (LSFM / STPT) or 2D histological slices. | Allen Mouse Brain Atlas (CCFv3) |
| **The Brain Globe Ecosystem** | Python API and Napari GUI. Minimal coding. Uses 3D volumetric (LSFM / STPT). | Allen Mouse Brain Atlas (CCFv3), Waxholm Rat, Princeton Mouse, Zebra Finch, and 20+ other species |
| **WholeBrain & SMART** | Python/Ilastik backend platform. Coding experience required. Uses 2D histological slice series with interactive checkpoints. | Allen Mouse Brain Atlas (CCFv3) |
| **DMC-BrainMap** | Unified multi-organism & multi-omic analysis; No coding required. Uses 2D histological slices. | Allen Mouse Brain Atlas (CFFv3), Waxholm Rat, Macaque Brain Atlas, |
| **BrainJ** | Uses FIJI ecosystem (plugin). No coding experience required. Automated serial 2D histological section reconstruction. | Allen Mouse Brain Atlas (CCFv3) |
| **SHARCQ Pipeline** | MATLAB-based, step-by-step GUI, Warps 2D histological sections to register to atlas. | Allen Mouse Brain Atlas (CCFv3) |
| **QUINT pipeline** | No coding required. GUI-based. Uses 3D volumetric (LSFM / STPT) or 2D histological slices. | CCFv3, Waxholm Atlas of the Spraque Dawley Rat (WHS), Kim Unified Adylt Mouse Brain Atlas, DeMBA, KimLabDev CCFv001), and potentially more (check source). |
| Atlas Registration Workflows |  |  |
| **QuickNII & DeepSlice** along with **QCAlign & MeshView** | No coding required. GUI-based. Uses 2D histological slices to reconstruct the 3D volume. | Allen Mouse Brain Atlas (CCFv3), Waxholm Rat Atlas, and Developmental Mouse Brain Atlas |
| **MADI3D (Virtual Fly Brain)** | Free GUI tool for Drosophila; desktop application. No coding. Uses 2D histological slices to reconstruct the 3D volume. | JRC2018, Ito2014, Jenett/JFRC2010, Court2018, Cardona/Janelia, Wood2018, JRC_FlyEM_Hemibrain |
| Cell Detection Workflows |  |  |
| **TRAILMAP** | Python-based. Coding experience required. 3D axonal projections & continuous fibers. Uses 3D volumetric datasets. | Allen Mouse Brain Atlas (CCFv3) |
| **Whole Mouse Brain 3D Cell Counting Pipeline** | R-based. Uses RStudio environment. Coding experience required. Uses 3D volumetric (LSFM / STPT) or 2D histological slices. | Allen Mouse Brain Atlas (CCFv3). Early postnatal Developmental Mouse Brain Atlas (epDevAtlas) |
| **CellMap** | Python-based. Requires python proficiency. Uses 3D volumetric datasets. | Allen Mouse Brain Atlas (CCFv3) |

Whole-brain imaging with cell detection mapped to a brain atlas produces complex datasets, inherently creating a post-processing chokepoint. To obtain accurate labeling, brain images need to be carefully aligned to a reference atlas (Puchades et al., 2025). Multiple interfaces, workflows, and pipelines have been established to align mechanical (sliced) vs optical (LSFM/STPT) sectioned images with reference atlases (Table 1). Some pipelines enable atlas alignment for a variety of species, including mouse, Drosophila, zebrafish, and macaque [(Yates et al., 2019, Jung et al., 2026), NeuroInfo; MBF Bioscience]. After atlas alignment, these workflows perform automated or semi-automated cell detection on the imaged markers for each aligned region. The cell detection outputs consist of cell count, density (cells/mm^3^), and region name, which are used for downstream analysis, interpretation, and visualization. Due to the high dimensionality and complexity of these outputs, parsing them for downstream analysis poses a significant challenge within the context of 3D brain architecture. The brain’s ontology is determined by the reference atlases used in these workflows. Of the available brain atlases, the mouse is commonly used and well-established, first by Paxinos and Franklin (Paxinos & Franklin, 2004) and more recently by the Allen Institute for Brain Science (AIBS). The AIBS provides standard 2D (ARA) and 3D (CCFv3) mouse brain reference atlases, both of which use the ARA ontology (Dong, 2008, Wang et al., 2020). Since the brain images are 3D-reconstructed, different brain depths correspond to specific regions that can be tracked through a hierarchical system. The AIBS’s ARA ontology takes advantage of this hierarchical system by starting at the “root” and extending to varying numbers of “leaves” in a hierarchical tree (Figure 1). This can be thought of as a parent-to-descendant representation of regions and subregions within a dendrogram; as you go further down the “family tree,” the depth increases. The root starts at Depth 0 and increases in depth level with each subregion and further division into sub-subregions.

**Figure 1.**
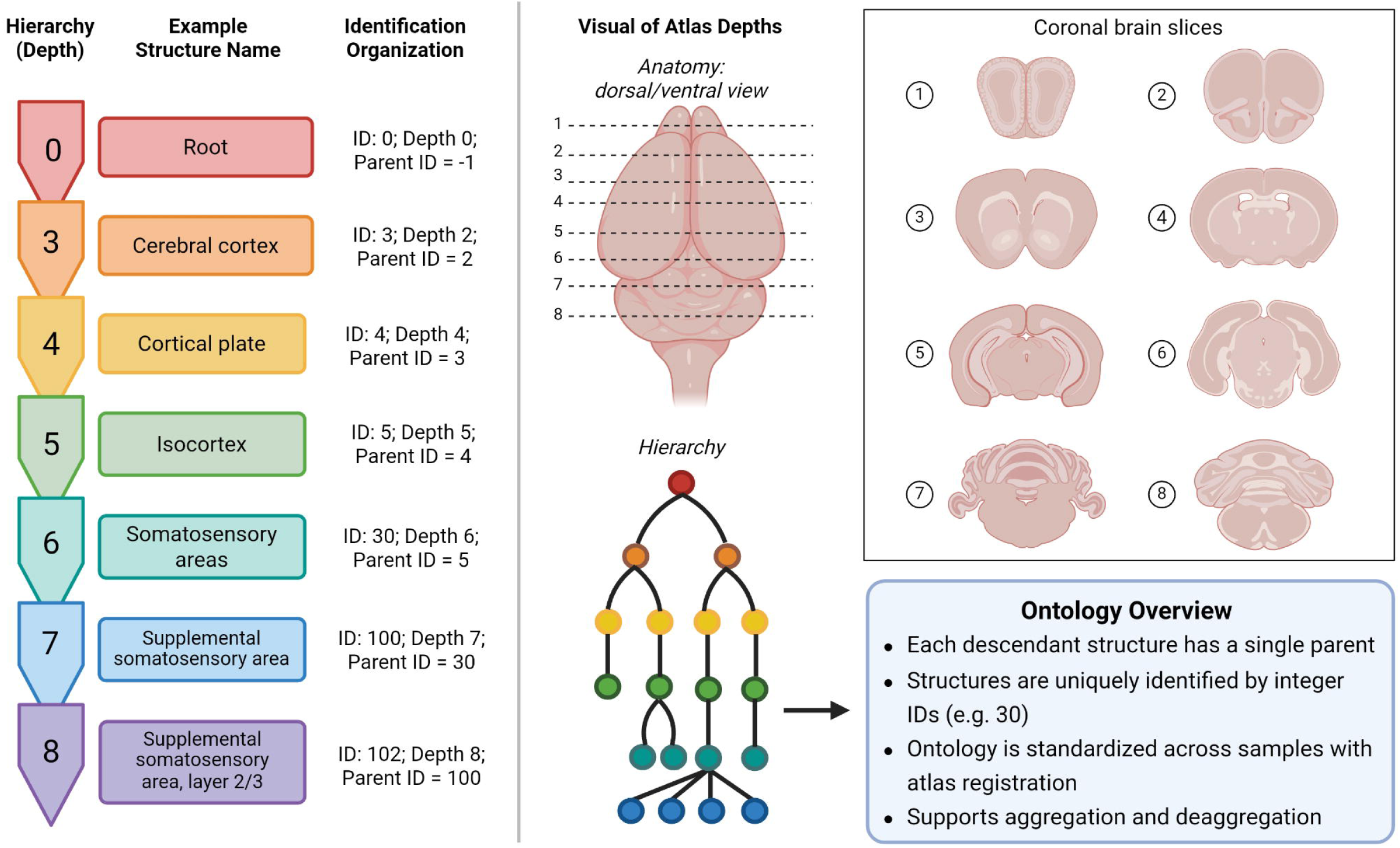
Atlas ontology overview. The diagram demonstrates the organization and naming scheme of 3D atlases such as CCFv3. The ontology follows a hierarchical structure from parent regions to descendant subregions. This hierarchical organization is how HERO refines whole-brain cell count datasets to select and rank the top-expressing regions. Created with BioRender.

Considering hierarchical organization in data interpretation from whole-brain cell detection datasets is especially important for mapping multi-scale granularity from macro-level (e.g., functional networks) systems to micro-regions (e.g., single-cell resolution), making it essential for contextualizing complex structural and functional data (Wang et al., 2019). Although hierarchical information provides more robust analysis of whole-brain datasets, utilizing this hierarchical structure complicates data interpretation, especially for those unfamiliar with nested ontologies. Whole-brain cell detection datasets often contain hundreds of registered regions, making direct interpretation difficult without some form of filtering or prioritization. Because of this complexity, methods for region selection can vary widely between studies, creating inconsistency and hindering rigor and reproducibility of results.

There are three main methods for approaching cell-detection datasets: 1) Data Reduction, 2) All-Region Screening, and 3) Network Analysis. Each approach identifies regions of interest with unique information that must be considered before selecting the desired path. The last two, all-region screening and network analysis, use complex, sophisticated mathematical models and theory to identify regions and relationships. All-region screening compares each atlas region between groups, usually using region-level cell counts or densities, and applying a multiple-comparison correction (Vasylieva et al., 2026). Significant regions are then selected and used for data interpretation and presentation. Network analysis with fixed tissues requires linking brain region expression patterns of activity markers, such as the immediate early genes cFos, Arc, and npas4, where higher marker expression indicates greater regional activity. These region-level activity data are then mapped and analyzed as networks, in which regions become nodes and relationship patterns become edges or graph features, similar to how functional network data from fMRI are treated (Hu et al., 2024; Rubinov & Sporns, 2010). The most common approach, especially among those newer to the field, is data reduction, which filters out regions for downstream statistics and data visualization. Within the data reduction approach, there are various methods for filtering these regions, such as by specific depth, density rankings, or biological interest (Chan et al., 2026, Jiang et al., 2026, Manjila et al., 2025, Patwardhan et al., 2025, Luchsinger et al., 2021). Consequently, reducing the dataset without accounting for anatomical hierarchy can distort biological interpretation. For example, multiple high-ranking subregions may belong to the same parent structure, creating the appearance of distributed brain-wide involvement when the signal may instead reflect enrichment within a single anatomically defined family. This issue is particularly relevant when users select the “top N” regions directly from deeper atlas levels, where fine anatomical subdivisions can dominate the ranked output and inflate the apparent importance of a single parent structure. A standardized, replicable approach to data reduction for whole-brain cell detection datasets is currently lacking, necessitating a more standardized workflow for the field. A hierarchy-aware strategy to cell-detection data reduction will robustly leverage this approach and standardize whole-brain analysis, further advancing biomedical research.

We developed HERO (Hierarchy-aware Expression Region Organization), a robust data reduction pipeline for whole-brain cell detection datasets. The HERO pipeline integrates a hierarchical organization system and accounts for laterality to refine and reduce data. HERO integrates atlas depth filtering, dynamic laterality handling, recursive parent-descendant organization, and iterative region balancing to identify key anatomical structures for downstream analysis. Rather than ranking all regions as independent units, HERO begins with broad parent structures (e.g., midbrain, cortical subplate) and then searches within their descendants to identify high-expressing descendant regions. This hierarchy-aware strategy reduces the risk of one highly subdivided parent structure inflating the final ranked list (i.e., amygdalar areas). It also improves upon fixed-depth selection by allowing regions without descendants at a target depth to remain eligible for inclusion in the final ranked list. HERO also preserves subject-level density values throughout the workflow, enabling both group-level and individual-level approaches. To improve interpretability, HERO incorporates hierarchy-guided collapse-and-refill logic, dense-ranking metrics, and customizable chord diagram visualizations to compare shared and group-specific expression patterns, while also providing users with this information in .xlsx file outputs. Additionally, HERO offers an optional focused-analysis feature that allows targeted interrogation of user-defined anatomical parent regions while maintaining the same hierarchy-aware framework.

To demonstrate the utility of this pipeline, we applied HERO to two newly generated mouse whole-brain afferent tracing datasets that replicate prior connectivity work from our group (Luchsinger et al., 2021), providing known afferent organization datasets to evaluate the workflow: one mapping inputs into the bed nucleus of the stria terminalis (BST), and the other tracing inputs to the insular cortex (insula). Both regions receive inputs from a wide variety of regions, yielding the kind of large complex cell detection datasets that HERO was designed to reduce into digestible summaries. Together, HERO provides a standardized and scalable framework for initial whole-brain cell detection analysis that improves interpretability and facilitates reproducible data reduction methods of atlas-organized datasets.

## 2 Methods

These methods provide a user-friendly preprocessing pipeline for raw cell-detection datasets from whole-brain imaging. HERO consists of five major steps: depth filtering, group-level averaging, hierarchical-guided region selection, overrepresentation correction, and visualization.

### 2.1 Methods for Acquiring Example Data

#### 2.1.1 Animals

All experiments were conducted in accordance with the National Institutes of Health Guide for the Care and Use of Laboratory Animals and approved by the Wake Forest University Institutional Animal Care and Use Committee. Six (2M/4F) adult C57BL/6J mice (The Jackson Laboratory; Bar Harbor, ME) were used. All animals were acclimated in standard group housing for 1 week upon arrival and handled for at least 1 week prior to an experiment. Animals were maintained on a 12h reverse light/dark cycle (lights on at 1800 h) and under controlled temperature (20-25°C) and humidity (30-50%) levels. Mice were given *ad libitum* access to food and water.

#### 2.1.2 Stereotaxic surgeries

Adult mice were anesthetized with isoflurane (3% initial dose, 1.5% maintenance dose) for intracranial adenosine-associated virus (AAV) injection surgeries. AAVrg-CAG-tdTomato (Addgene #59462) virus was injected with 300 nl at a rate of 50 nl/min into the right BST for three female mice (from the bregma: A/P +0.20, M/L -2.13, D/V -3.88, 15° angle) or the right mid-insular cortex (from bregma: A/P +0.02; M/L -3.80; D/V −4.20; 0° angle) for one female and 2 male mice. All mice recovered for at least 4 weeks before tissue extraction. Mice underwent transcardial perfusions with 4% paraformaldehyde (PFA) with 0.01mg/ml heparin. Brains were extracted and incubated in 4% PFA for 24h before further processing.

### 2.2 Whole-Brain Light-Sheet Preparation, Imaging, and Cell Detection

#### 2.2.1 Whole-brain tissue clearing, index matching, and imaging

PFA-fixed samples were preserved with SHIELD reagents (LifeCanvas Technologies, LCT) according to the manufacturer’s instructions (Park et al., 2018) and as previously described (Luchsinger et al., 2021). Samples were delipidated using LCT Clear+ delipidation reagents. Samples were incubated in 50% Index matching solution (EasyIndex, RI = 1.52, LCT) overnight at 37°C, followed by a 1-day incubation in 100% EasyIndex for refractive index matching. Samples were mounted in 2% ultra-low melt agarose made with EasyIndex and placed into the indexed matched immersion medium on the microscope and incubated overnight. The samples were imaged on a LifeCanvas SmartSPIM light-sheet axially swept microscope using a 3.6x objective (0.2 NA) (LCT). All images were acquired at 4 um increments and subsequently de-striped and stitched for analysis.

#### 2.2.2 Whole-brain atlas registration and cell detection

Images were registered to the Allen Brain Atlas (AIBS: https://portal.brain-map.org/) using a two-phase semi-automated process in SmartAnalytics (LCT). For each brain, the autofluorescence channel was registered to an averaged autofluorescence reference atlas generated from previously aligned specimens by LCT. The atlas alignment/registration workflow applied a sequence of rigid, affine, and b-spline warping algorithms (SimpleElastix: https://simpleelastix.github.io/; *more information on warping*: https://lifecanvastech.com/how-to-use-the-allen-brain-atlas/). This multistep approach ensures both global and fine-scale anatomical correspondence between each sample and the atlas.

Automated cell detection data was generated by SmartAnalytics using a custom convolutional neural network built with the TensorFlow Python package. Cell detection consists of two sequential models. First, a fully convolutional detection network (https://arxiv.org/abs/1605.06211v1) based on a U-Net architecture (https://arxiv.org/abs/1505.04597v1) identifies candidate cell locations across the volume. Second, these candidates are evaluated by a classifier network employing a ResNet (https://arxiv.org/abs/1512.03385v1) architecture to determine whether each location represents a true-positive cell. After detection, cell coordinates were added onto the Allen Brain Atlas (AIBS CCFv3) using the previously computed registration, enabling quantification of cell counts within atlas-defined anatomical regions. The AIBS CCFv3 ontology used for the mouse structure order can be found in the Supplementary Table S1.

Cell detection outputs are formatted as CSV files with columns for region ID, region name, acronym, parent ID, depth, count, volume, and density. This information is required to apply the HERO pipeline. 3D videos showing the injection site and afferent projections are included in Supplementary Materials for the BST (Supplementary Video 1) and the insula (Supplementary Video 2).

For other workflows that conduct atlas registration and cell detection, see Table 1. Resources on scalable brain atlases can be found at the Human Brain Project (HBP: https://www.humanbrainproject.eu/en/science-development/focus-areas/brain-atlases/#whs) and the Scalable Brain Atlas (SBA: https://scalablebrainatlas.incf.org/).

### 2.3 HERO Methods

Whole-brain expression data is exported from SmartAnalytics as a region-level CSV file containing region ID, region name, acronym, parent ID, depth, count, volume, and density. SmartAnalytics output follows the hierarchical atlas structure of the Allen Brain Atlas, and HERO is designed to preserve anatomical organization while identifying high-density regions across samples and experimental groups using a hierarchy-aware framework. HERO provides a hierarchy-aware alternative to standard ranked lists by starting from broad parent regions (e.g., midbrain, cortical plate) and identifying the highest-expression descendants within those anatomical lineages. This reduces parent-region overrepresentation while preserving meaningful high-expressing regions that may not extend to the same atlas depth. HERO consists of five major steps: depth filtering, group-level averaging, hierarchy-guided region selection, overrepresentation correction, and visualization (Figure 2). The pipeline is written in R therefore, R must be installed before using the pipeline (install R: https://cran.wustl.edu/). HERO can be cloned from GitHub: (https://github.com/avashipman/HERO).

**Figure 2.**
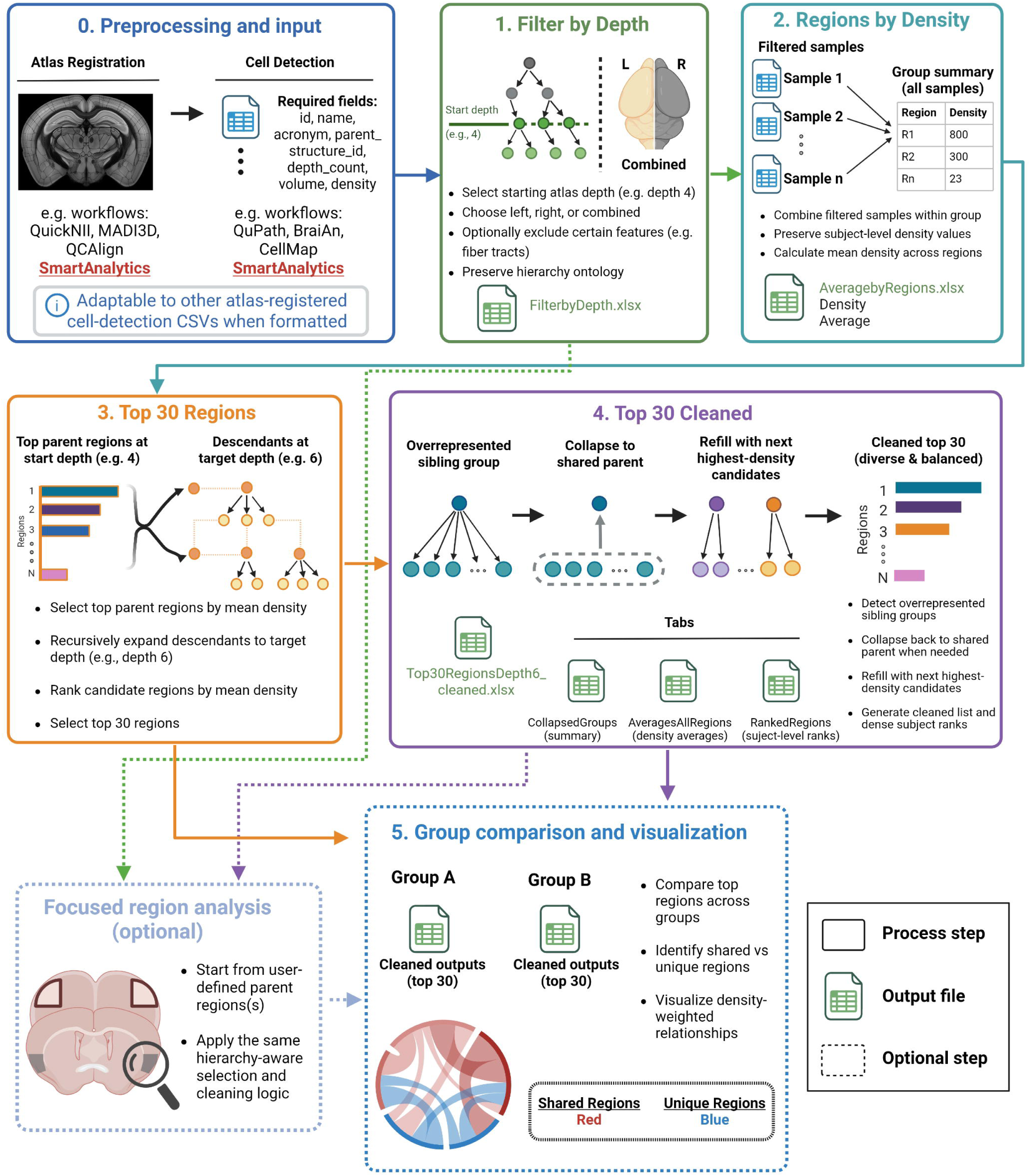
HERO workflow. Workflow and the corresponding output files generated for each HERO step. The process includes optional steps such as focused region analysis, group comparison, and visualization. HERO is a *plug-and-chug* pipeline in R that refines whole-brain cell count datasets for interpretation. Created with BioRender.

#### 2.3.1 Input data and atlas hierarchy (Figure 2, Step 0.)

Each sample is represented by a CSV file containing region-specific measurements. Required fields include the region ID (*id*), region name (*name*), region acronym (*acronym*), hierarchical anatomical organization (*parent_structure_id*), regional depth (*depth*), cell count (*count*), volume (*volume (mm^3)*), and density (*density (cells/mm^3)*). HERO’s *‘parent_structure_id’* parameter is used to reconstruct the atlas hierarchy, where each descendant region references the *‘id’* value of its anatomical parent. The *‘depth’* variable is used to define the anatomical resolution of the analysis, with broader brain regions occurring at lower depths and more specific subregions occurring at higher depths. Table 1 provides a list of example open-source pipelines for atlas registration and cell detection of 2D histological slices or 3D volumetric datasets. This list does not include commercial software packages such as Bruker (IMARIS), MBF Biosciences (Neuroinfo, BrightSLICE), and LifeCanvas Technologies (SmartAnalytics). When inputting cell-detection datasets from workflows other than SmartAnalytics into HERO, they need to be formatted with the above-mentioned syntax (as italicized above) for each of the required fields. If these criteria are met, HERO can be applied to any whole-brain cell detection dataset.

#### 2.3.2 Step 1: depth filtering and laterality (Figure 2, Step 1.)

The first step, implemented by the ‘FilterbyDepth’ function, filters each sample-level CSV file to a user-defined atlas depth to identify the parent structures for the hierarchy-aware analysis part of the HERO pipeline. Beyond the hierarchical system of the ARA ontology, the CCFv3 atlas incorporates laterality by distinguishing left and right hemispheres, adding another layer of complexity. It also divides the brain into gray matter, fiber tracts, and ventricular systems. The gray matter is then subdivided into three large regions: “cerebrum”, “brain stem”, and “cerebellum”. The CCFv3 then categorizes 12 “major divisions” of the gray matter: isocortex, olfactory areas, hippocampal formation, cortical subplate, striatum, pallidum, thalamus, hypothalamus, midbrain, pons, medulla, and cerebellar areas (Wang et al., 2019, Wang et al., 2020). These 12 major divisions are largely represented at Depth 4. Since HERO uses broad atlas structures as parent anchors for identifying the highest-expressing descendants, Depth 4 is recommended as the standard starting depth to select parent structures. However, the function allows the user to dynamically specify any starting depth level.

The function also provides dynamic laterality handling if the cell-detection datasets contain it. For single hemisphere analysis, only regions from the selected hemisphere are retained. For combined analyses, left and right homologous regions are merged by summing expression counts and regional volumes separately, followed by recalculation of density:

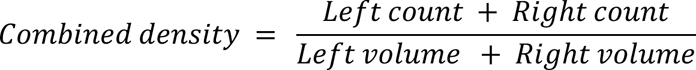

Combined densities thus reflect true values rather than an average of hemispheric densities. Cleaned region names and acronyms are generated by removing laterality-specific prefixes or suffixes, while preserving the original anatomical information needed for filtering. For datasets that already combine laterality prior to HERO, select the *combined* option in the ‘laterality’ parameter.

Each sample fed into the HERO pipeline produces a filtered Excel file containing the selected regions, density values, region IDs, depth information, and hierarchy metadata.

#### 2.3.3 Step 2: group-level density averaging (Figure 2, Step 2.)

The second step, ‘RegionsbyDensity’, combines the filtered Excel files from *Step 1* across all samples in a group. For each anatomical region, the function calculates the mean density across samples while preserving individual subject-level density values. HERO primarily uses density to rank regions, as this metric normalizes volume, allowing regions with different volumes to be compared. Differences in cell count and density are compared in Supplementary Figures 1C-D and 2C-D.

The output file, ‘AverageRegions.xlsx’, contains two tabs: The ‘Density’ tab stores individual sample density values, with samples as rows and brain regions as columns. This tab can be directly pasted into visualization tools such as GraphPad to visualize top regions from the starting depth (e.g., Depth 4). The ‘Averages’ tab contains each region’s mean density, atlas ID, and depth, sorted from highest to lowest density. This file serves as the starting point for HERO’s selection of highly expressed regions.

#### 2.3.4 Step 3: hierarchy-guided region selection (Figure 2, Step 3.)

The third step, ‘Top30Regions’, identifies the highest-density parent regions from ‘AveragedRegions.xlsx’ and proceeds through the atlas hierarchy to select the top 30 expressing descendants, or the number specified by the user. The function selects the top parent regions at the starting depth (produced by *Steps 1 & 2*), then uses the parent-descendant relationships in the original SmartAnalytics CSV files to recursively identify descendants until the target depth is reached. HERO’s default target depth is Depth 6, as it provides sufficient specificity to include key regions such as the BST and the insula. However, the user can specify their target depth. For each candidate’s descendant region, density values are extracted for every sample. Mean density was then calculated across samples, and the top 30 regions are selected. This first-pass selection can be viewed in the ‘Top30RegionsDepth6_.xlsx’ output file.

#### 2.3.5 Step 4: Correction for parent overrepresentation (Figure 2, Step 4.)

Since some anatomical structures, such as hypothalamic or cerebellar regions, contain many detailed regions at certain depths, highly granular parents can dominate the top 30 list. To address this, the ‘Top30Cleaned’ function applies a hierarchy-aware correction for data refinement.

The function identifies three or more selected Depth 6 descendant regions that originate from the same previous depth parent (e.g., Depth 5). If this occurs, the descendant regions collapse back into the shared parent regions. The parent density is then used to represent that anatomical group. After collapsing, the function refills the remaining slots with the next-highest-density candidate regions from the same hierarchy-defined candidate pool. Refilling prioritizes Depth 6 regions, then Depth 5 regions, and uses the original starting depth only as a final fallback. This iterative collapse-and-refill process continues until the final list contains 30 regions without overrepresentation from any single parent structure. The removed descendant regions and the number of iterations taken for each collapsed region are also documented in the final output file. The ‘collapse_threshold’ parameter is defaulted to two but can be adjusted by the user.

The cleaned Excel file contains several tabs. The ‘Densities’ tab contains the final cleaned region list with the mean density, acronym, and depth. The ‘AveragesAllRegions’ tab documents the full candidate pool considered during selection, including regions excluded during collapse. The ‘CollapsedGroups’ tab records all parent-descendant collapses. The ‘RankedRegions’ tab ranks all candidate regions by density within each subject using dense ranking and reports the group-level mean and median density. The tabs offer the user flexibility in data interpretation preferences and quality checking of the HERO pipeline. For example, if a user does not want to collapse descendant regions, the ‘AveragesAllRegions’ tab shows the uncleaned top 30. Another example is if an expected region of interest does not show up in the cleaned Top 30 list. Since collapsed regions are shown in the ‘CollapsedGroups’ tab, the user can reference this tab to determine if the region of interest was collapsed and decide whether to include it in the user’s interpretation. Overall, this flexibility makes HERO a powerful and dynamic tool for data interpretation.

#### 2.3.6 Step 5 (Optional): focused region analysis and group visualization (Figure 2, optional)

##### 2.3.6.1 Focused region analysis

A separate, and optional, focused-analysis function is available for targeted investigation of specific anatomical parent regions. Rather than beginning with the highest-density parent regions, this function allows the user to select one or more starting regions at a specified depth. The function then identifies all descendants of the specified parent region(s) until the target depth is reached. Candidate descendants are ranked by mean density across samples, and the same overrepresentation correction and refill strategy used in ‘Top30Cleaned’ is applied. This enables focused investigation of anatomically defined regions while maintaining the same hierarchy-aware selection logic.

##### 2.3.6.2 Group comparison and visualization

To compare expression patterns between experimental groups, cleaned top-region outputs can be visualized using chord diagrams with HERO’s “make_density_chord_parentgroups’ function. Each group’s cleaned output file is used as input. To avoid overrepresentation by one group, the visualization function selects the top 30 regions by default, which can be user-adjusted, within each group separately, and plots only those selected group-region relationships.

The chord diagram shows ribbon-like chords connecting brain regions to experimental groups. The ribbon thickness is proportional to density, where thicker ribbons represent higher densities. Regions in the top 30 that appear in more than one group are labeled as shared, with corresponding ribbons colored red. In comparison, ribbons for regions present in only one group are colored with the corresponding group specified by the user. Region ordering was based on the ‘ParentRegion’ field from the ‘AveragesAllRegions’ tab, allowing anatomically related regions to cluster together visually. This visualization does not show statistically different or similar regions; rather, it displays how regions are prioritized differently and similarly in cell detection. HERO’s visualization function provides a first-pass impression of how groups differ in their cell detection, easily summarizing both group-specific and shared high-density expression patterns while preserving hierarchy-aware anatomical organization.

## 3 Results

### 3.1 Applications of HERO

To demonstrate the utility of HERO as a downstream data reduction and interpretation workflow, we applied the pipeline to two separate whole-brain retrograde tracing cell-detection datasets: 1) the bed nucleus of the stria terminalis (BST; Supplementary Video 1) and 2) the insular cortex (insula; Supplementary Video 2). These two regions have afferents spanning multiple anatomical systems, requiring a workflow like HERO to reduce their large atlas-registered outputs to interpretable, exploratory regional summaries. A list of the brain regions and their acronyms selected by HERO is found in Supplementary Table 2.

Cell density is the primary metric used to rank regions in HERO as it normalizes volume. Cell count measurements can provide useful information, but they are not as compatible for comparing regions with different volumes. Ranked lists of cell density and counts were compared and showed that rankings depended on which measurement was used (Supplementary Figures 1C-D and 2C-D). For example, MO is a large region that ranked first in the count list. However, when looking at density, it was not included, showing that it had a low enough concentration (cells/mm3) of marked cells to not make the top 30 list for the BST dataset (Supplementary Figure 1C-D). This example demonstrates how volume can prioritize larger regions and potentially overlook important structures with smaller volumes. Altogether, density is the most optimal measurement to use in HERO as it normalizes volume when comparing regions across depths and granularity in ranked lists.

For the example datasets, HERO analysis was performed using a starting atlas depth of 4 (depth_level = 4) and a target depth of 6 (target_depth = 6). Left and right hemispheres were combined, and the 30 highest-density parent regions were selected (top_n_parents = 30). Candidate descendants were ranked with the top 30 (top_n_final = 30). For refinement, a collapse threshold of two (collapse_threshold = 2) was applied.

#### 3.1.1 HERO applied to BST whole-brain afferent tracing dataset

Using the HERO workflow on the BST retrograde tracing cell-detection dataset, the top 30 parent regions were first selected starting at atlas Depth 4, and their descendants at Depth 6 were ranked from highest to lowest BST-projecting cell density. The resulting top 30 list identified the highest-density candidate input regions in Depth 6 to the BST prior to hierarchy-aware collapsing (Figure 3D). This initial ranked output provides a broad exploratory overview of regions with high fluorescent cell density and is consistent to conventional depth data-reduction methods (Supplementary Figure 1B-C). Since a large number of closely related descendant structures reside in Depth 6, HERO identified dominant structures projecting to the BST such as the isocortex, hypothalamus, and the periventricular zone (Figure 3E).

**Figure 3.**
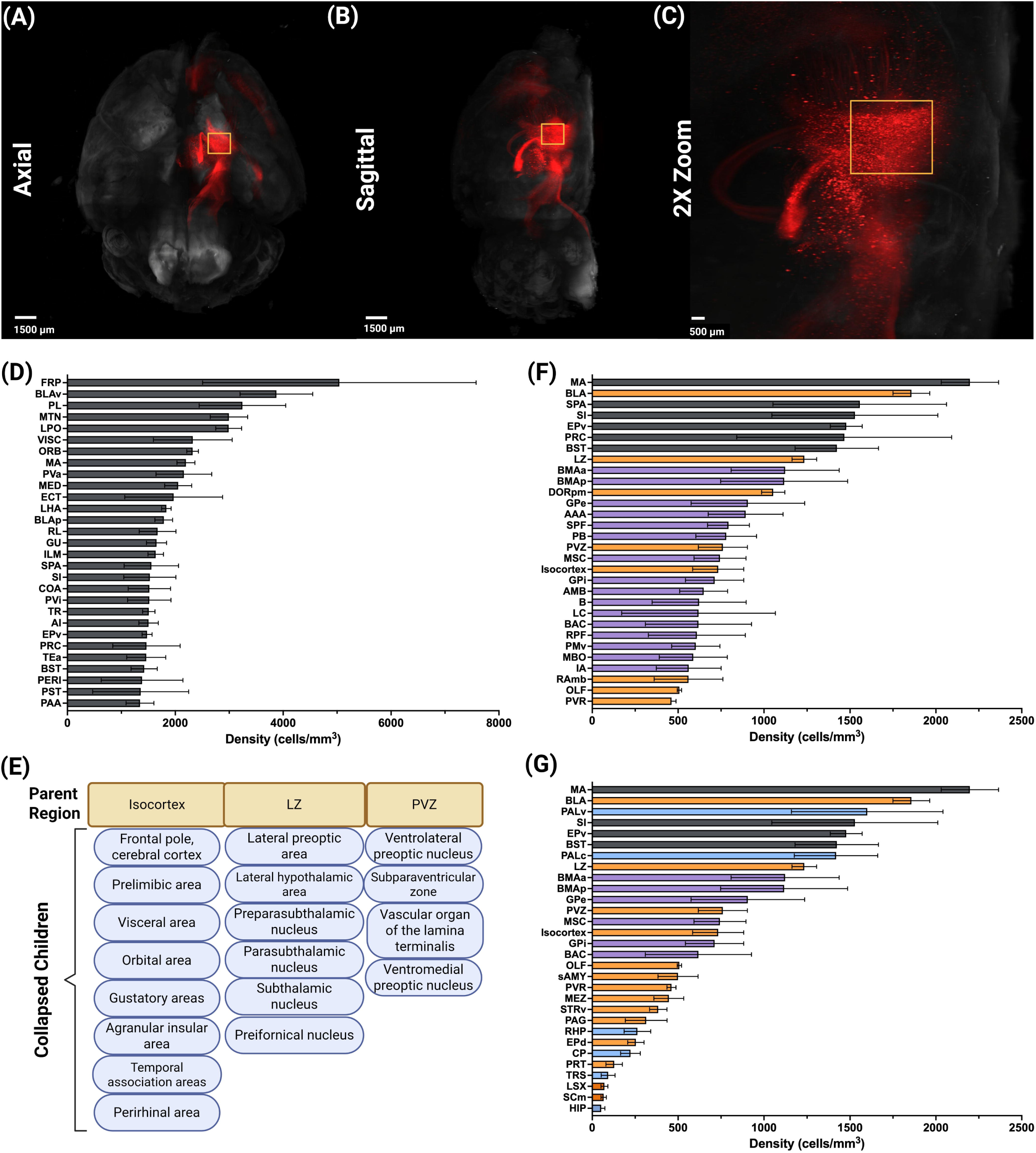
Applying HERO to a whole-brain BST afferent mapping dataset. Representative image of afferent projections to the right BST in **(A)** axial, **(B)** sagittal, and **(C)** 2X zoom on the injection site (AAVrg.TdTomato). The right BST injection site is highlighted in the yellow box for each panel. **(D)** HERO’s uncleaned data reduction approach showing the top 30 regions expressing BST projection cells from Depth 6. **(E)** Example of selected descendants that met the criteria for collapsing into their parent regions. **(F)** Cleaned top 30 regions using HERO’s ‘collapse descendants’ feature. Gray bars signify regions that were also identified in the standard top 30. Orange bars signify parent regions of collapsed descendants. Purple bars signify unique regions in the top 30 from the cleaned version. **(G)** Focus analysis using selected Depth 4 parent regions. Blue bars signify unique regions identified in the focused version, but not the cleaned version.

To identify regions in Depth 6 that are not from these dominant areas, the hierarchy-aware cleaning step was applied to the initial top 30-ranked outputs with a collapse threshold of 2 (Figure 3F). Overrepresented descendant regions were identified (e.g., Figure 3E), collapsed to their shared parent structure when appropriate, and the remaining positions in the ranked list were filled with the next-highest-density candidate regions (Figure 3F). This produced a cleaned top 30 output that retained high-density BST-projecting regions while reducing redundancy among closely related anatomical descendants. With the cleaned output, other regions, such as the RAmb, OLF, and RPF, were identified (Figure 3F).

Comparison of the uncleaned and cleaned ranked lists demonstrates the value of the HERO refinement step. The uncleaned output highlights the highest density structures. In contrast, the cleaned output provides a more balanced anatomical summary by preventing a single parent region or a closely related group of descendant regions from dominating the final list. HERO, therefore, distinguished between dense labeling of a subdivided anatomical group and broader distribution projection patterns across the brain.

To rank regions from larger parent areas of interest, we used the optional focus HERO function, “Top30FocusedCleaned” and selected the following parent structures: pallidum, hypothalamus, cortical plate, cortical subplate, and hippocampal formation (Figure 3G). The focused analysis revealed more memory-related structures such as the HIP and RHP. Predictably, the pallidum and cortical plate remain the densest regions containing BST-projecting cells.

#### 3.1.2 HERO applied to whole-brain insula afferent dataset

Mapping insula afferents is a strong example of the overrepresentation of closely related groups of descendant regions that dominate ranked lists. Using the exploratory HERO workflow, the top 30 parent regions were first selected starting at atlas Depth 4, and their descendants at Depth 6 were ranked from highest to lowest insula-projecting cell density. The resulting top 30 list identified the highest-density candidate input regions in Depth 6 to the insula prior to hierarchy-aware collapsing (Figure 4D). This list is also consistent to other data-reduction depth methods, retaining dense regions with the hierarchical framework (Supplementary Figure 2B-C). Most regions in the ranked list come from autonomic systems such as the pons (p-mot) and include regions from the isocortex and hypothalamus (Figure 4E). However, when the hierarchy-aware cleaning step in the HERO pipeline is applied, the list is populated with regions involved in reward, memory, and attention signaling, such as the globus pallidus, hippocampus, and basal forebrain (Figure 4F), providing a more dynamic and balanced regional expression summary.

**Figure 4.**
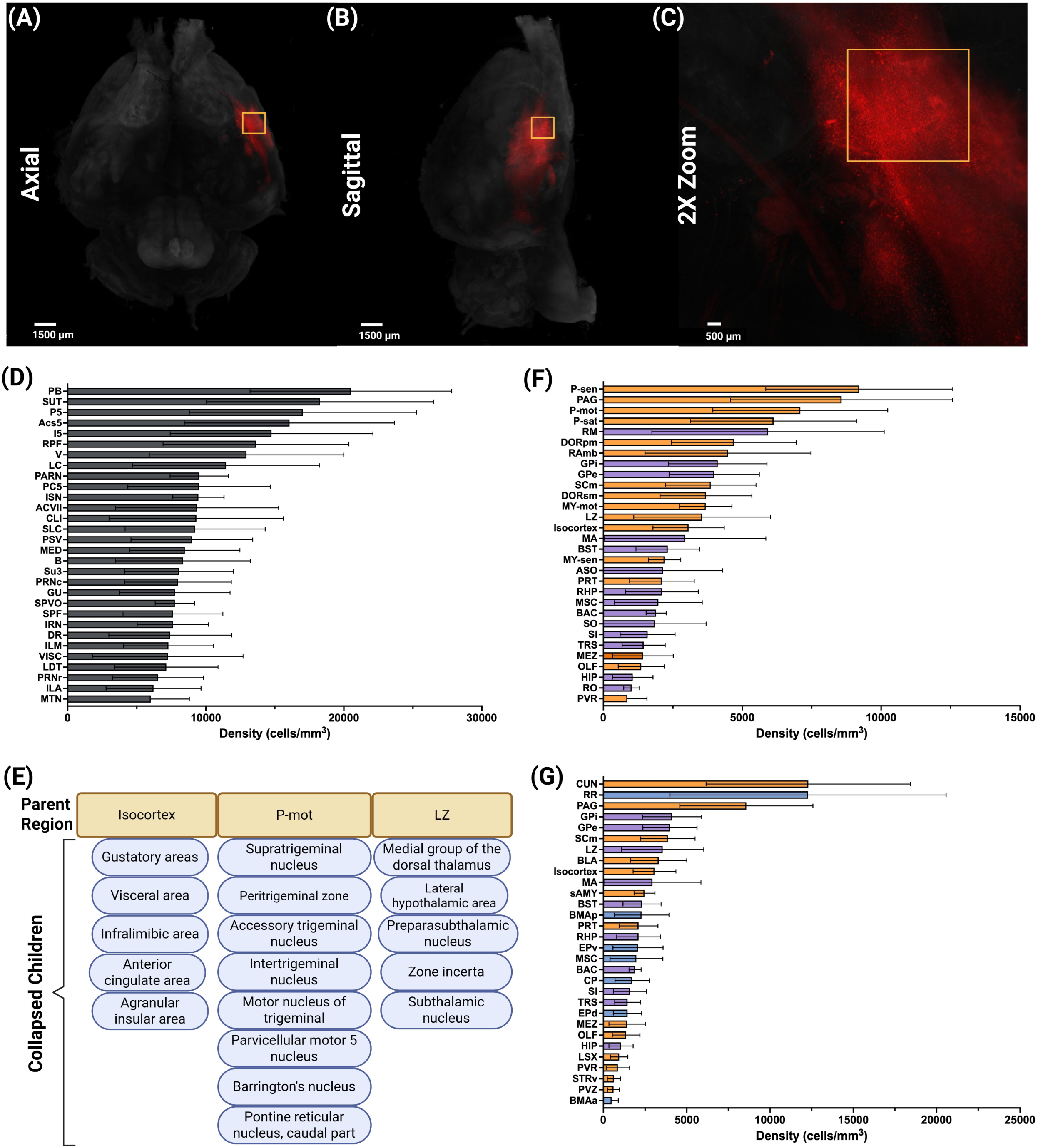
Applying HERO to a whole-brain insula afferent mapping dataset. Representative image of afferent projections to the right insula in **(A)** axial, **(B)** sagittal, and **(C)** 2X zoom on the injection site (AAVrg.TdTomato). The right insula injection site is highlighted in the yellow box for each panel. **(D)** HERO’s uncleaned data reduction approach showing the top 30 regions expressing insula projection cells from Depth 6. **(E)** Example of selected descendants that met the criteria for collapsing into their parent regions. **(F)** Cleaned top 30 regions using HERO’s ‘collapse descendants’ feature. Gray bars signify regions that were also identified in the standard top 30. Orange bars signify parent regions of collapsed descendants. Purple bars signify unique regions in the top 30 from the cleaned version. **(G)** Focus analysis using selected Depth 4 parent regions. Blue bars signify unique regions identified in the focused version, but not the cleaned version.

To rank regions from larger parent areas of interest, we used the optional focus HERO function, Top30FocusedCleaned and selected parent structures strongly associated with reward, stress, and emotion processing: pallidum, hypothalamus, cortical plate, cortical subplate, and hippocampal formation (Figure 4G). Within these selected parent structures, the top descendants included the hippocampus, amygdala, the lateral septum, and the isocortex. This list provides evidence of enriched regions projecting to the insula within these preselected areas.

#### 3.1.3 Comparing HERO-identified top BST-projecting regions to top insula-projecting regions

Comparing top regions expressing BST- and insula-projecting cells can provide valuable information about each region’s afferents and shared projection patterns. The uncleaned HERO analysis reveals that the top regions projecting to the BST and insula are distinct instead of shared (Figure 5A). However, plotting the cleaned HERO analysis reveals substantial overlap of top afferent regions (Figure 5B). Thus, the cleaned function can add valuable information with a broad anatomical coverage perspective that other data reduction methods that do not use hierarchical organization would miss. These shared regions include SI, MA, LZ, PVR, DORpm, and the globus pallidus, which are loosely linked to hormone secretion (e.g., oxytocin and vasopressin) and to regulating internal stress responses (Goel et al., 2025).

**Figure 5.**
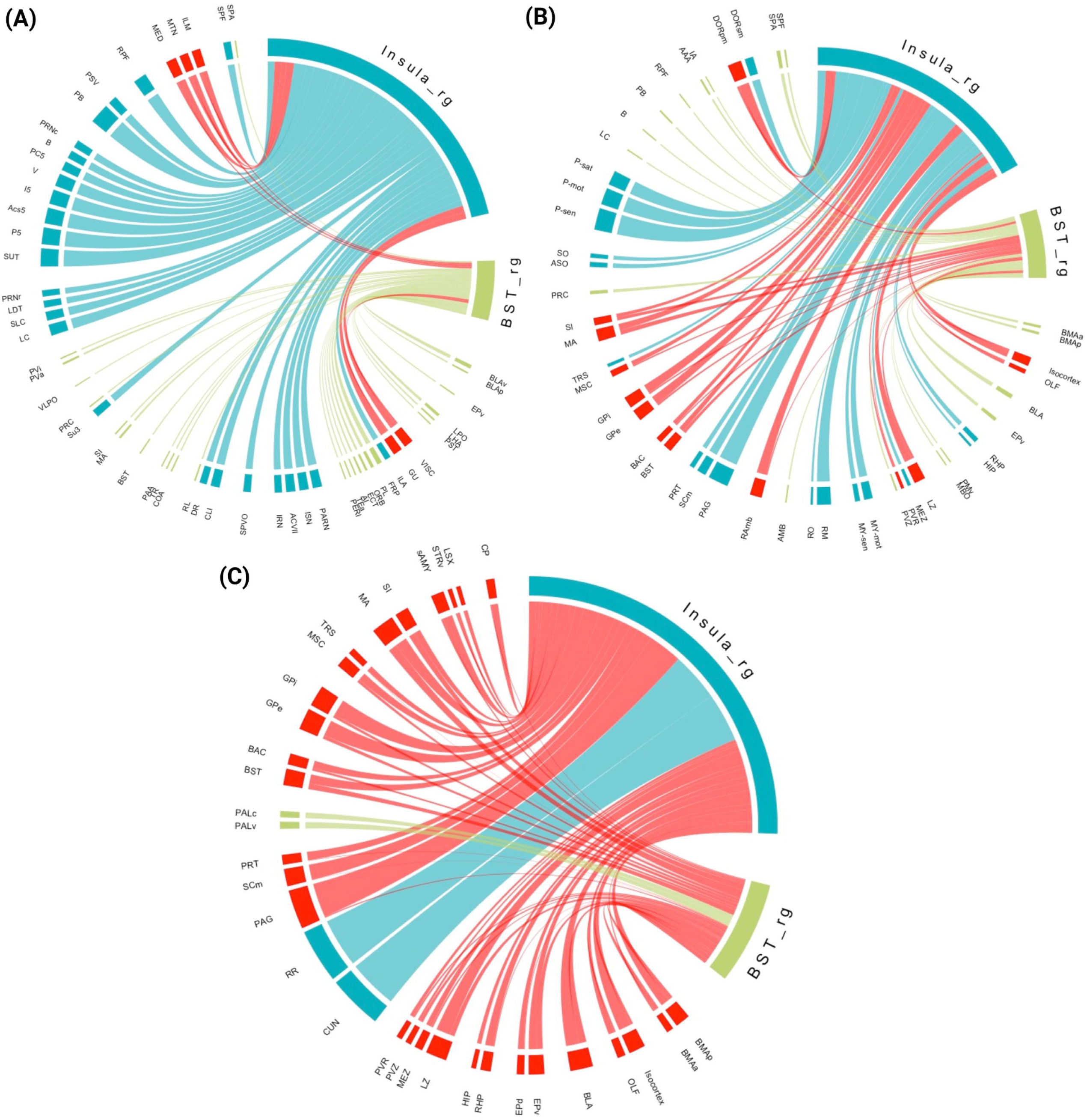
Comparing top regions projecting to the BST and the insula. **(A)** Chord diagram visualizing differences between the top 30 insula- and BST-projecting regions. Blue ribbons represent the top 30 afferent regions identified in the insula but not the BST. Green ribbons represent the top 30 afferent regions identified in the BST but not in the insula, and red ribbons indicate shared regions ranked in the top 30 for both the insula and BST afferents. **(B)** Chord diagram visualizing differences between the top 30 insula- and BST-projecting regions from the cleaned rank list. **(C)** Chord diagram visualizing differences between the top 30 insula- and BST-projecting regions from the focused rank list.

Since most of the regions shared between the insula and BST were associated with autonomic functions, we used the HERO-focused analysis feature to select parent regions involved in higher somatic control and to identify the top expressing descendants in Depth 6: pallidum, hypothalamus, cortical plate, cortical subplate, and hippocampal formation. The focused analysis revealed more overlap of regions, especially in the amygdalar areas (e.g., BMAa, BLA, BST), thalamic areas (e.g., SI, MA, LZ, PVZ), the isocortex, and the globus pallidus (Figure 5C).

### 3.2 Customizable Features of HERO and Sensitivity Analysis

To appeal to a wide range of users, HERO’s parameters for ranking region densities are flexible and depend on the perspective the user needs. Due to this flexibility, parameters such as the collapse threshold will influence which regions are represented in the final ranked lists (Supplementary Figure 3). To assess the sensitivity of the output to the adjustable HERO parameters, a similarity index test for the top 30 lists was calculated using the Jaccard similarity index, defined as the size of the intersection divided by the size of the union of the selected top 30 lists, where 1 indicates exact similarity, and 0 indicates no similarity. Similarity index compared the cleaned top 30 lists at different threshold levels to our chosen threshold 2, which is an aggressive collapsing threshold (Supplementary Figure 3B). For the insula dataset, thresholds 1 and 2 produced identical cleaned selections. However, increasing the threshold to 3 dramatically changed the selected regions, with the similarity decreasing to 0.154. Threshold 4 further declined to 0.034. Thresholds 4-6 produced identical final ranked selections to each other. The BST dataset, on the other hand, showed more gradual changes across thresholds. Compared to threshold 2, similarity was 0.25 at threshold 1, 0.33 at threshold 3, and 0.15 at threshold 6. Both experiments retained the anatomical representation of all 30 original high-density regions, meaning the ancestry of the originally selected regions remained in the cleaned top 30 outputs. However, the insula dataset replaced all original top 30 regions with their ancestors, whereas the BST dataset retained the four original top 30-ranked regions (Figure 3–4). When comparing the similarity index between the uncleaned and cleaned top 30 lists at different thresholds, both the insula and BST datasets show low similarity indices (Supplementary Figure 3C). A lack of similarity between uncleaned and cleaned ranked lists is an expected consequence of replacing descendants with parents. The uncleaned lists emphasize region-specific high-density structures, while the cleaned lists emphasize broader, anatomically nonoverlapping regions. How broad the regions can become depends on the collapsing threshold. For example, less aggressive thresholds 4-6 provide a different balance between high-density specificity and anatomical coverage (Supplementary Figure 3C).

Since the cleaned and uncleaned lists have differing regions but retain anatomical representation, the preservation of uncleaned density information was calculated by density attenuation between each collapsed descendant and their parent structure 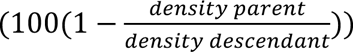. Higher-density attenuation results in a greater loss of original density information. Attenuation calculations for each collapsed descendant (Supplementary Figure 3D) further demonstrate that aggressive collapsing lowers the density used to represent originally highly expressing descendants. For the insula dataset, all 30 originally selected high-density regions were replaced by broader anatomical ancestors, confirming retention of anatomical coverage. The median attenuation in the insula dataset was 48.5% (Supplementary Figure 3D). Among the 34 collapsed children, 16 exhibited greater than 50% attenuation, using the collapse threshold of 2. For the BST dataset, the median attenuation was 51.7%. Four original high-density regions were retained in the cleaned outputs, while 26 were represented through ancestors. Among the total 30 collapsed descendants, 16 exhibited greater than 50% density attenuation. Density attenuation calculations are included in all cleaned outputs, flagging users of collapsed children with greater than 50% density attenuation.

Overall, HERO enables users to explore large whole-brain cell-detection datasets through flexible parameters that allow multiple perspectives for qualitatively examining the data. Users should test different collapse thresholds to determine the sensitivity of the results to collapsing. Since the parameters affect the cleaned ranked outputs, users should report their parameters when using HERO for reproducibility.

## 4 Discussion

Mouse mesoscale imaging has transformed the ability to quantify cellular expression patterns across intact brains. When paired with atlas-based registration, these datasets offer an unprecedented opportunity to identify distributed marker patterns across experimental groups. However, interpreting and refining these outputs remains a major challenge. Atlas-registered datasets are inherently hierarchical, with each region embedded within parent-descendant anatomical relationships that vary in depth, scale, and biological specificity. As a result, simple ranking approaches that select regions solely by density, count, or regions of interest can unintentionally overrepresent anatomically related regions, obscure broader circuit-level patterns, and have biased interpretations. The Hierarchy-aware Expression Region Organization (HERO) pipeline was developed to address this analytical gap by providing a workflow for selecting, refining, ranking, and visualizing whole-brain datasets. HERO gives users a defined path from analyzed 3D volumetric or reconstructed histological slice datasets to biologically meaningful interpretations, promoting exploratory analysis and lowering the barrier to incorporating mesoscale imaging.

### 4.1 HERO’s Advantages Over Other Approaches for Analyzing Whole-Brain Cell Detection Datasets

The sheer magnitude of whole-brain cell detection datasets underscores the need for a standardized approach to data reduction or region prioritization. There are three main approaches for cell detection datasets: 1) Data Reduction, 2) All-Region Screening, and 3) Network Analysis. Each has its own purpose, but the most common, especially for those newer to the field, is data reduction. Data reduction allows users to select regions with a specific criterion for downstream statistics and visualization. However, these criteria vary across the field, such as using targeted regions of interest (e.g., reward-related regions) or using targeted depth (e.g., Depth 6). As these approaches have led to insightful discoveries (Chan et al., 2026, Patwardhan et al., 2025), they differ widely from each other while also obscuring crucial information that whole-brain imaging enables, such as other depth levels and parent-descendant region groupings. HERO addresses these issues by incorporating hierarchical organization into ranked cell detection regions, providing detailed intermediate outputs, and supporting metrics. A central advantage of HERO is that it preserves anatomical context during data reduction and improves upon this standard approach by incorporating depth-based filtering and parent-descendant relationship tracking into the analytical workflow (Figure 2). This allows users to define the level of anatomical resolution most appropriate for their experimental question while reducing redundancy among closely related regions. In doing so, HERO supports a more robust and well-rounded interpretation of whole-brain patterns by retaining biologically meaningful anatomical relationships rather than treating each atlas entry as an independent structure.

The workflow also improves transparency in how regions are selected and prioritized. Many whole-brain imaging studies employ a range of effective region-selection strategies, including top-ranked lists (as in previous studies from our lab (Luchsinger et al., 2021), expert-driven manual curation (Franceschini et al., 2023), and investigator-defined regions of interest (Chan et al., 2026), each well-suited to the questions these studies set out to address. The field currently lacks an accessible standardized format for presenting whole brain datasets. HERO addresses this by generating a guided, customizable workflow with traceable intermediate outputs, including depth-filtered datasets, averaged regional density tables, ranked region outputs, and cleaned top-region lists. These outputs allow users to explore datasets beyond a single ranked list, examine how parent-descendant relationships influence region selection, and assess cell marker patterns at both individual-subject and group levels. For novice whole brain imaging researchers, this structured approach provides an accessible starting point for navigating complex anatomical hierarchies, understanding how analytical parameters influence results, and identifying biologically meaningful patterns. This transparency is particularly important for method development, where reproducibility, interpretability, and an open science mentality are central criteria for advancing science.

### 4.2 Applying HERO to Example Data

#### 4.2.1 Top 30 ranked lists

Here, we generated and analyzed two whole-brain cell-detection datasets to demonstrate HERO’s utility with retrograde tracing of two brain areas our lab has expertise in: The BST and insula. The various functions within HERO provided vital information on dense projecting regions that dominate the list, showed obscured regions unique to the insula and BST, and revealed key dense regions shared between the insula and BST. Conventional rankings at Depth 6 are compared with HERO’s uncleaned top 30 regions expressing BST-projecting cells (Supplementary Figure 1B-C), showing that HERO’s uncleaned rankings provide a standard overview of enriched areas at the chosen depths that project to the BST. This also verifies that HERO can effectively isolate top regions that align with previous rodent literature (Beyeler and Dabrowska, 2020, Guerrero-Hreins et al., 2026, Sun et al., 2023, Luchsinger et al., 2021). The top regions, such as amygdalar areas, hypothalamic areas, and the temporal association area, are also consistent with human literature, showing HERO’s ability to capture the translatability in rodent BST tracing studies (Weller and Smith, 1982, Avery et al., 2014). Outside the dominant areas (Figure 3E), the Top30Cleaned function collapsed around 20 regions from the major areas found in Figure 3E, resulting in a more anatomically distributed list of top regions (Figure 3F). Instead of most regions being associated with homeostasis and the limbic system, we find a more diverse set of top-expressing regions involved in emotional, motivational, and stress-response processes (Cui et al., 2017, Goel et al., 2025, Jones, 2020).

For regions expressing insula-projecting cells, the conventional top 30 regions are comparable to HERO’s uncleaned top 30 list (Supplementary Figure 1B-C) and are consistent with other insula tracing studies (Gehrlach et al., 2020, Gogolla, 2017, Luchsinger et al., 2021). To compare to human studies, HERO’s top 30 regions, such as intra-insular regions, basal nuclei, and somatosensory regions, were also among human insular connectivity literature (Augustine, 1996, Uddin et al., 2017). When the Top30Cleaned HERO function was applied (Figure 4F), additional regions associated with memory, emotion, sensory integration, and neuroendocrine secretion and homeostasis were included (Goel et al., 2025, LeDoux et al., 1985, Zhu et al., 2025, Choi et al., 2023, Anacker et al., 2016). Altogether, HERO’s refinement step notably listed other key regions that project to the insula that would’ve otherwise been obscured with traditional ranking methods.

#### 4.2.2 HERO provides an extensive overview of whole-brain cell detection datasets

The visualization components of HERO support interpretation of biological group comparisons by converting dense region-level outputs into more accessible summaries. HERO creates chord diagrams to compare projecting regions for BST and the insula (Figure 5). First, the uncleaned top 30 regions were visualized between the two (Figure 5A). Initially, the chord diagram shows many unique ribbons, emphasizing that the raw ranked list indicates these two regions have separate inputs. With further inspection of the chord diagram, the separation is mainly due to strong projections from regions in the Pons to the insula, which the BST lacks. On the other hand, the BST contains more amygdalar regions than the insula. However, when collapsing these granular regions and prioritizing anatomical coverage, the refined ranked list shows an increase in the number of shared regions in the top 30 list for the BST and the insula (Figure 5B). These shared top regions do not provide evidence of overlapping BST and insula afferent neuronal populations but rather demonstrate similar top projection patterns with a broader anatomical perspective. The shared regions include the isocortex, amygdalar areas, thalamic, and hypothalamic regions, to highlight a few. Most of these shared regions come from stress, neuroendocrine regulation, and sensory relay areas (Goel et al., 2025, Choi et al., 2023, Anacker et al., 2016).

Another feature of HERO is the focused function, which allows the user to isolate parent regions of interest and select the most highly expressing descendants (Figure 3G and 4G). In the focus shared chord diagram (Figure 5C), top-expressing regions shared between the BST and the insula include the BLA, HIP, and striatum regions. Recent studies have been exploring the insula and BST in relation to these regions while investigating reward- and stress-related behaviors, showing the relevance of these shared regions in the field (Salin et al., 2027, Sun et al., 2024, Luyck et al., 2020, Zhou et al., 2026, Essoh et al., 2026).

Overall, application of HERO demonstrates effective discernment of relevant enriched regions for insula and BST projections. Additionally, with HERO’s refinement logic for collapsing closely related descendants within a hierarchical framework, previously obscured enriched regions were revealed, providing interpretable information in large cell-detection datasets. HERO also enabled focused visualization of top regions from selected parent regions of interest. Lastly, HERO provided visual comparison between insula and BST projecting regions to provide a broad perspective on the similarities and differences in enriched projecting regions. The different formats produced by the HERO workflow enable users to thoroughly explore whole-brain cell detection datasets from a range of perspectives. The uncleaned version of the hierarchy ranking reveals regions that can clump together to form larger areas that dominate expression. The cleaned version collapses these dominating larger areas to include more evenly distributed regions in the ranked list. The focused version enables users to identify important subregions in larger brain areas of interest (i.e., reward-related areas). The chord diagrams effectively visualize overarching group comparisons.

### 4.3 Adaptability of HERO Across Whole-Brain Atlas-Registered Datasets

HERO is broadly applicable across different types of expression-based whole-brain datasets. Although retrograde tracing is used in our example dataset, the workflow is not limited to this method. HERO can be applied to datasets generated from any marker that produces region-level counts, volume, and density following atlas registration. This may include other markers for neuronal activation, cell identity, projection labeling, receptor expression, disease pathology, or genetically labeled cell populations. Therefore, the pipeline is not restricted to a single biological question or experimental design. Rather, HERO provides a flexible analytical framework for interpreting any whole-brain dataset in which regional expression or labeling patterns are summarized within a hierarchical anatomical ontology.

Although HERO was developed using SmartAnalytics outputs, the conceptual framework can be adapted to other atlas-registered datasets (Table 1) if the necessary input structure is preserved. In its current implementation, HERO requires the input CSV files to contain the same types of information and formatting provided by SmartAnalytics, including columns containing region ID (*id*), region name (*name*), region acronym (*acronym*), hierarchical anatomical organization (*parent_structure_id*), regional depth (*depth*), count (*count*), volume (*volume (mm^3)*), and density (*density (cells/mm^3)*). Common workflows that conduct both atlas registration and segmentation for cell counts such as ClearMap2, QUINT, and ABBA+BraiAn contain this information and may require additional steps to obtain a compatible format for HERO’s input structure. ClearMap2 labels hierarchical information as graph order (o); ABBA+BraiAn contains parent IDs from image analysis with QuPath, while QUINT retains this information in its QCAlign plugin. Cell detection datasets from proprietary software such as IMARIS, Neuroinfo, and BrightSLICE could potentially be used with HERO, but their compatibility has not been verified. If users are unable to find the hierarchical anatomical organization in their whole-brain cell-detection datasets, the reference atlases should have the hierarchical order that users can manually add to be compatible with HERO (e.g., AIBS CCFv3 structure order in Supplementary Table S1). After generating whole-brain cell detection datasets using workflows such as those outlined above and in Table 1, and formatting them to HERO’s input requirements, the platform can be applied for data reduction. We encourage users to test HERO with other whole-brain datasets beyond SmartAnalytics. HERO is not restricted to the per-region metric it operates on. Although density is used here, the workflow applies equally to any measure summarized by brain region, including mean fluorescence intensity from bulk labeling or axonal metrics derived from segmentation tools such as TRAILMAP. Structural measures, such as regional axon volume, could therefore be organized and compared through HERO once quantified per region, though we have not tested this directly.

### 4.4 Caveats, Considerations, and Future Development

With all the benefits HERO offers, it is not intended to replace detailed statistical testing or region-specific follow-up analyses. Rather, it serves as an organizational and interpretive framework that identifies meaningful patterns in large anatomical datasets prior to downstream statistical analysis and validation.

HERO operates on the assumption that proper quality control has confirmed the accuracy of upstream imaging, registration, segmentation, atlas registration, and marker counting prior to its use. Therefore, proper quality checks should be conducted during the preprocess to generate the most accurate HERO outputs.

While using HERO, parameter settings such as depth and collapse threshold will influence the ranked outputs (Supplementary Figure 3), requiring users to report on the chosen parameters for transparency and reproducibility. Throughout the workflow, HERO provides quality-check measures that users should follow. Each HERO output provides ranked lists by subject for all candidate regions beyond the top 30 list so users can qualitatively identify outliers and determine if the average density of a region is driven by an individual subregion. A unique feature of HERO is the CleanedTop30 outputs, which use the collapse-and-refill process. It requires a collapse threshold that is defined by the user, and we have shown that the ranked lists are sensitive to the chosen value (Supplementary Figure 3B-C). A more aggressive collapse threshold of 2-3 preserves anatomical representation in high-density regions. However, the parent regions frequently exhibit substantially lower densities than their high-expression descendants (Supplementary Figure 3D). More mild collapse thresholds, such as 4-6, retain higher levels of signal attenuation because they have more descendants that represent the overall parent region. This indicates that more aggressive hierarchical aggregation prioritizes anatomical representation over localized high-density signals, thereby reducing redundancy within a single larger area but lacking regional specificity.

Utilizing the uncleaned and cleaned outputs as complementary ranked lists provides users a well-rounded foundation of the patterns seen in their cell-detection datasets. The user should check the CollapsedGroups tab in the output to have a record of the collapse-and-refill process for each iteration, density attenuation values, and flagging of collapsed descendants. The density attenuation values and flagging further reveal how descendants are represented within their ancestry and whether a high-density signal may be locally concentrated within a descendant structure, a unique perspective that HERO provides over traditional data reduction methods. This will facilitate determining which threshold is preferred and reporting specific parameters used for transparency and reproducibility. If the user solely wants an unfiltered list of all the candidate regions from the starting depth to the target depth, the AveragesAllRegions tab and the uncleaned list provide that. Furthermore, overlap validation is conducted to verify that refill regions are unique to the collapsed ancestry in the SelectionReport tab. By generating ranked tables and group-level visualizations, HERO helps users identify dominant regional patterns while preserving information about shared and unique expression profiles across groups. HERO was designed to curate datasets to a level that is accessible across from novice neuroanatomist to experts, serving as a learning tool while encouraging hypothesis generation, simplifying figure generation, and facilitating effective science communication.

Additionally, HERO requires sample CSVs to have columns with the following information: region ID (*id*), region name (*name*), region acronym (*acronym*), hierarchical anatomical organization (*parent_structure_id*), regional depth (*depth*), cell count (*count*), volume (*volume (mm^3)*), and density (*density (cells/mm^3)*). Since HERO’s main approach relies on the hierarchical organization of brain region labels, parent region IDs are necessary. Most open-source whole-brain analysis workflows do not have this information in their initial data outputs and may require additional steps. If users utilize the CCRFv3 atlas for registration, parent region IDs can be found in the Supplementary Material (Supplementary Table S1) and easily implemented with HERO. However, other atlases could be harder to track and find parent region IDs if not already provided by pre-HERO pipelines.

HERO is limited by the availability of brain atlases for other species, as many lack standardized, high-quality reference atlases comparable to the mouse brain for registering their whole-brain samples. Only a handful of pipelines work with other species atlases (see Table 1), underscoring the need for the field to create standardized atlases for a wide range of species. While HERO reduces overrepresentation among related regions, choices such as depth level, ranking metric, and collapse threshold remain user-defined parameters. These decisions should be reported clearly to ensure reproducibility across studies.

Future development of HERO will increase its accessibility and analytical power. One important direction is the development of a graphical user interface (GUI) that would allow users to run the workflow, adjust parameters, inspect intermediate outputs, and generate visualizations without writing code. HERO will also be optimized to support additional input formats, allowing broader compatibility within the workflow. Another important future direction is incorporating statistical analysis. Recent work by Kentaro Ishii and colleagues uses hierarchy-aware statistics for whole-brain cell-detection datasets with their BRANCH workflow in their preprint, demonstrating the power of considering anatomical relationships in these datasets and a shift in the field towards this direction (Ishii et al., 2025). Furthermore, since whole-brain expression datasets can capture distributed activity patterns with markers such as cFos, future versions of HERO will be extended to examine relationships among regions, identify coordinated expression networks, or integrate anatomical hierarchy with functional clustering. Such advances would move the pipeline beyond data reduction to include all-region screening and network analysis.

### 4.5 Conclusion

HERO provides a practical, user-friendly workflow for reducing whole-brain cell detection datasets while respecting the hierarchical structure of anatomical ontologies. By combining depth-aware filtering, region ranking, parent-descendant refinement, and visualization, HERO addresses a common bottleneck in whole-brain imaging analysis: transforming large atlas-registered datasets into biologically meaningful and reproducible interpretations. HERO improves upon current data reduction approaches by mitigating misleading overrepresentation of related anatomical regions and providing a more transparent starting point for exploratory analysis. The density attenuation values and flagging in the cleaned outputs further reveal how the hierarchical framework behaves, emphasizing the importance of descendant-parent relationships, a unique feature of HERO compared with traditional data reduction methods. Through customizable parameters, hierarchy-aware region selection, and transparent data refinement, HERO allows users to examine their datasets from multiple anatomical perspectives rather than relying on a single ranked list. Overall, HERO advances the accessibility, interpretability, and standardization of analyzing whole-brain cell detection datasets, providing a foundation for more informative biological discovery from brain-wide imaging.

## Supporting information

Suppelmental Material

## 5 Conflict of Interest

On behalf of all authors, the corresponding author states that there is no conflict of interest.

## 6 Author Contributions

ALS: Conceptualization, Investigation, Methodology, Software, Writing – original draft, Visualization, Validation. SWC: Writing – review & editing, Investigation, Supervision, Conceptualization.

## 7 Funding

This manuscript was supported by the National Institute of Health grant T32AA007565 (A.L.S.) and the Whitehall grant 2023-12-071 (S.W.C.).

## 8 Acknowledgments

We want to thank Jincy Little for her technical assistance with the brain sample preparation and Tatiyana Kubrynski, M.S., for testing and giving vital feedback on the HERO pipeline development. HERO was refactored, debugged, and optimized using ChatGPT (OpenAI). All code has been reviewed and tested by the authors. Figures 1–2 were created with BioRender. The Supplementary Videos 1 and 2 were created in Python with AI generated code (Gemini).

## 11 Data Availability Statement

The datasets used for this study can be found in the HERO GitHub repository https://github.com/avashipman/HERO.

## Notes

https://github.com/avashipman/HERO

### Competing Interest Statement

The authors have declared no competing interest.

### Summary of Updates

We have included additional Supplementary Materials (Supplemental Figures 1-3 and Videos 1-2) in the manuscript that 1) compare conventional top-n rankings with HEROs lists, 2) include in-depth validation tests, and 3) provide visualizations of density values of our whole-brain data in the form of coronal heat maps and 3D videos. The description of this supplementary information has been updated and is referenced in the Methods, Results, and Discussion. Figures 3 and 4 were revised.

