## Supplementary material for "HERO: A hierarchy-aware analysis pipeline for reducing and refining whole-brain atlas-mapped cellular datasets": Suppelmental Material

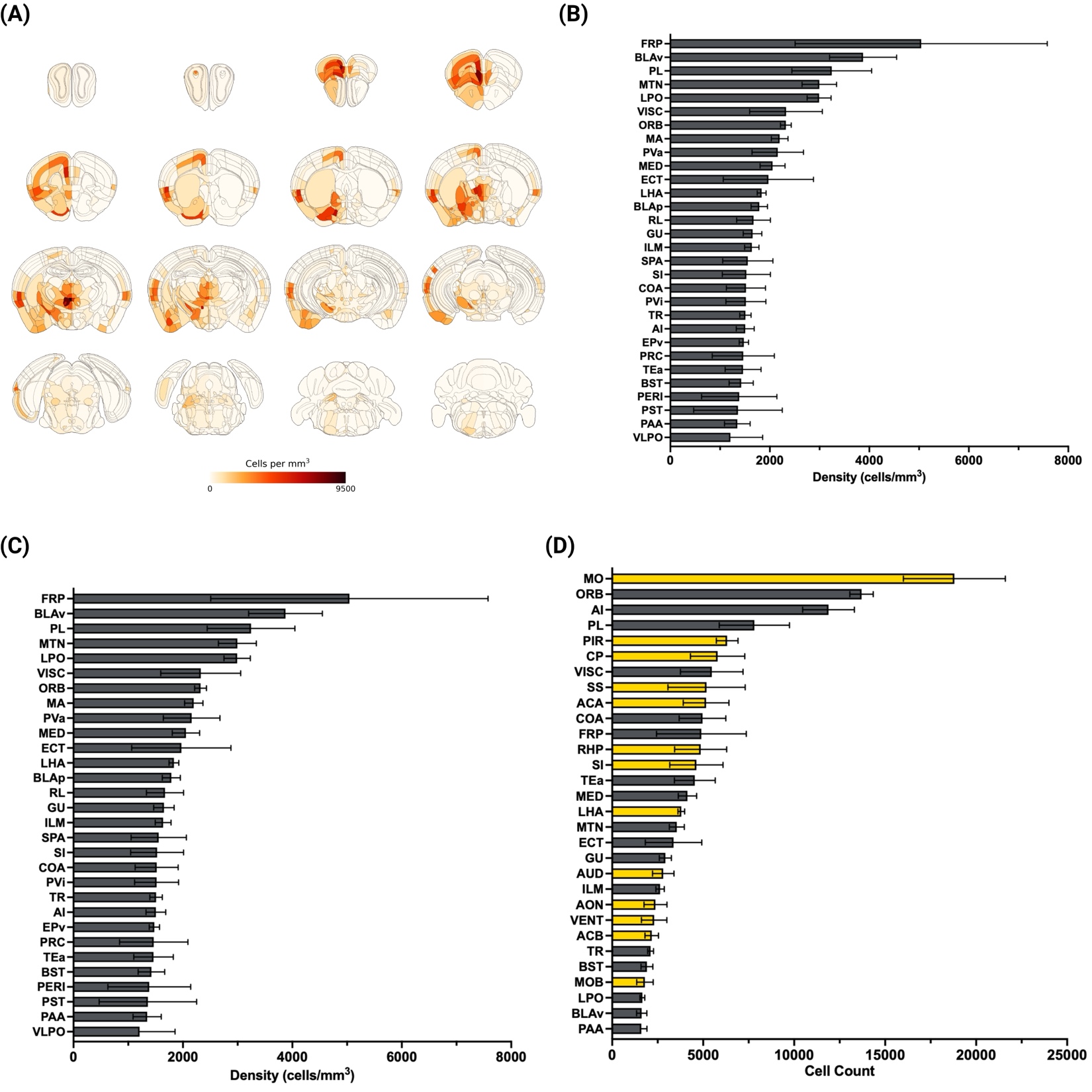

**Supplementary Figure 1**. **BST anatomical representations of density and comparisons of conventional depth filtering, HERO’s depth filtering, and density compared to cell counts**. **(A)** Representation of cell density on a composite whole-brain heatmap using matplotlib and BrainGlobe’s brainglobe-heatmap (Claudi et al., 2024). **(B)** Conventional Depth 6 ranking approach showing top 30 regions expressing BST projection cells. **(C)** HERO’s uncleaned top 30 highest density list at target Depth 6. **(D)** Top 30 highest cell count list at target Depth 6. Yellow bars signify unique regions ranked by using cell counts.

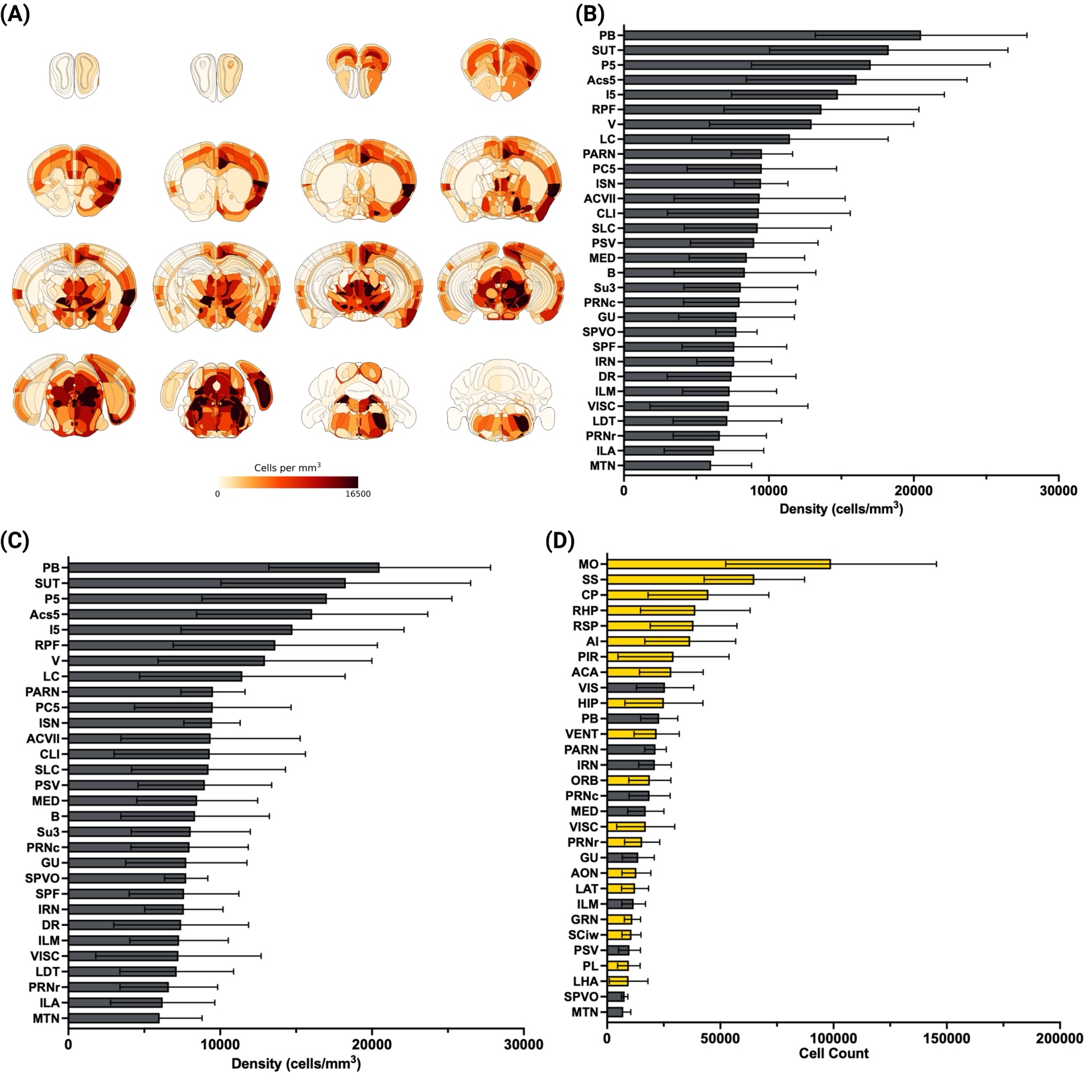

**Supplementary Figure 2**. **Insula anatomical representations of density and comparisons of conventional depth filtering, HERO’s depth filtering, and density compared to cell count**. **(A)** Representation of cell density on a composite whole-brain heatmap using matplotlib and BrainGlobe’s brainglobe-heatmap (Claudi et al., 2024). **(B)** Conventional Depth 6 ranking approach showing top 30 regions expressing insula projection cells. **(C)** HERO’s uncleaned top 30 highest density list at target Depth 6. **(D)** Top 30 highest cell count list at target Depth 6. Yellow bars signify unique regions ranked by using cell counts.

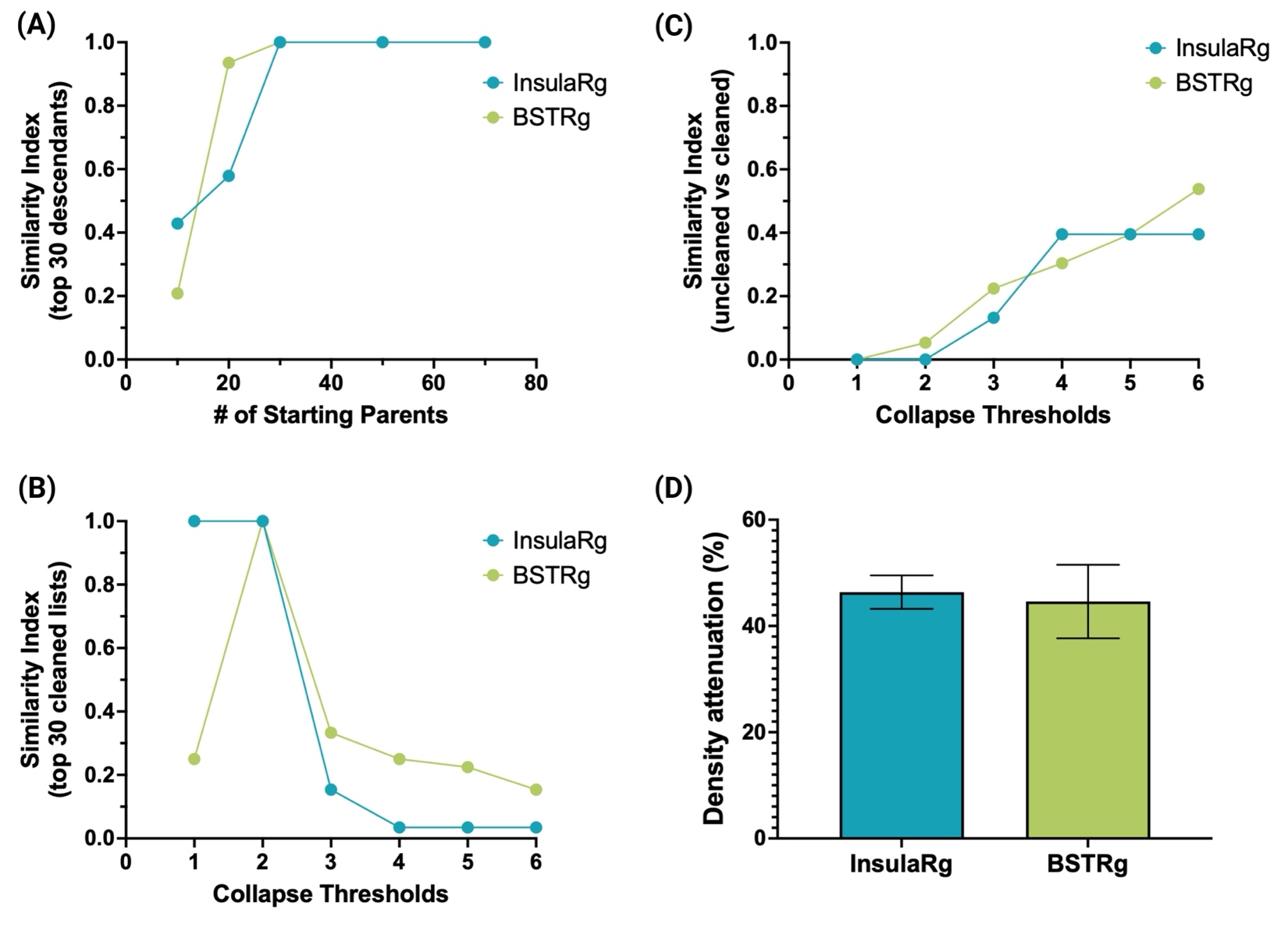

**Supplementary Figure 3**. **Sensitivity tests for changing HERO parameters**. **(A)** Similarity index comparing the starting number of parent regions for downstream Depth 6 ranking of descendants. **(B)** Similarity index comparing top 30 HERO cleaned outputs across thresholds using threshold 2 as the standard comparison. **(C)** Similarity index comparing uncleaned and cleaned HERO outputs across thresholds. **(D)** Density attenuation of each collapsed descendant from uncleaned to cleaned HERO outputs for each dataset when collapsing with threshold 2.

**Supplementary Video 1. Video of labeled afferents from the BST**. Light sheet imaging of AAVrg.tdTomato in a cleared whole mouse brain. Gray outlines the brain from the 488 nm autofluorescence. Yellow highlights the injection site which contains the largest contiguous region above an intensity threshold (4,000 arbitrary pixel units), overlaid so it remains visible from all angles. Display ranges are 250-6,000 au, gamma 0.6, and the scale bar is 2mm. Videos were made in Python with the source code available on GitHub (<https://github.com/avashipman/HERO/tree/main/RenderBrainVideo>).

**Supplementary Video 2. Video of labeled afferents from the insula.** Light sheet imaging of AAVrg.tdTomato in a cleared whole mouse brain. Gray outlines the brain from the 488 nm autofluorescence. Yellow highlights the injection site which contains the largest contiguous region above an intensity threshold (12,000 arbitrary pixel units), overlaid so it remains visible from all angles. Display ranges are 250-12,000 au, gamma 0.6, and the scale bar is 2mm. Videos were made in Python with the source code available on GitHub (<https://github.com/avashipman/HERO/tree/main/RenderBrainVideo>).

**Supplementary Table 2.** CCFv3 nomenclature and abbreviations of highlighted brain regions referred to in the example data

| **Abbreviation** | **Region nomenclature** |
| --- | --- |
| ACVII | Accessory facial motor nucleus |
| ASO | Accessory supraoptic group |
| Acs5 | Accessory trigeminal nucleus |
| AI | Agranular insular area |
| AAA | Anterior amygdalar area |
| ACA | Anterior cingulate area |
| AON | Anterior olfactory nucleus |
| AUD | Auditory areas |
| B | Barrington's nucleus |
| BLA | Basolateral amygdalar nucleus |
| BLAp | Basolateral amygdalar nucleus, posterior part |
| BLAv | Basolateral amygdalar nucleus, ventral part |
| BST | Bed nuclei of the stria terminalis |
| BAC | Bed nucleus of the anterior commissure |
| CP | Caudoputamen |
| CLI | Central linear nucleus raphe |
| COA | Cortical amygdalar area |
| CUN | Cuneiform nucleus |
| DR | Dorsal nucleus raphe |
| ECT | Ectorhinal area |
| EPd | Endopiriform nucleus, dorsal part |
| EPv | Endopiriform nucleus, ventral part |
| FRP | Frontal pole, cerebral cortex |
| GRN | Gigantocellular reticular nucleus |
| GPe | Globus pallidus, external segment |
| GPi | Globus pallidus, internal segment |
| GU | Gustatory areas |
| HIP | Hippocampal region |
| LZ | Hypothalamic lateral zone |
| MEZ | Hypothalamic medial zone |
| ISN | Inferior salivatory nucleus |
| ILA | Infralimbic area |
| IA | Intercalated amygdalar nucleus |
| IRN | Intermediate reticular nucleus |
| I5 | Intertrigeminal nucleus |
| ILM | Intralaminar nuclei of the dorsal thalamus |
| Isocortex | Isocortex |
| LAT | Lateral group of the dorsal thalamus |
| LHA | Lateral hypothalamic area |
| LPO | Lateral preoptic area |
| LSX | Lateral septal complex |
| LDT | Laterodorsal tegmental nucleus |
| LC | Locus ceruleus |
| MA | Magnocellular nucleus |
| MBO | Mammillary body |
| MED | Medial group of the dorsal thalamus |
| MSC | Medial septal complex |
| MY-mot | Medulla, motor related |
| MY-sen | Medulla, sensory related |
| RAmb | Midbrain raphe nuclei |
| RR | Midbrain reticular nucleus, retrorubral area |
| MTN | Midline group of the dorsal thalamus |
| V | Motor nucleus of trigeminal |
| AMB | Nucleus ambiguus |
| RM | Nucleus raphe magnus |
| RO | Nucleus raphe obscurus |
| OLF | Olfactory areas |
| ORB | Orbital area |
| PALv | Pallidum, ventral region |
| PB | Parabrachial nucleus |
| PC5 | Parvicellular motor 5 nucleus |
| PARN | Parvicellular reticular nucleus |
| PAG | Periaqueductal gray |
| PERI | Perirhinal area |
| P5 | Peritrigeminal zone |
| PVa | Periventricular hypothalamic nucleus, anterior part |
| PVi | Periventricular hypothalamic nucleus, intermediate part |
| PVR | Periventricular region |
| PVZ | Periventricular zone |
| PIR | Piriform area |
| PAA | Piriform-amygdalar area |
| P-sat | Pons, behavioral state related |
| P-mot | Pons, motor related |
| P-sen | Pons, sensory related |
| PRNr | Pontine reticular nucleus |
| PRNc | Pontine reticular nucleus, caudal part |
| TR | Postpiriform transition area |
| PRC | Precommissural nucleus |
| PL | Prelimbic area |
| PST | Preparasubthalamic nucleus |
| PRT | Pretectal region |
| PSV | Principal sensory nucleus of the trigeminal |
| RHP | Retrohippocampal region |
| RPF | Retroparafascicular nucleus |
| RSP | Retrosplenial area |
| RL | Rostral linear nucleus raphe |
| MO | Somatomotor areas |
| SS | Somatosensory areas |
| SPVO | Spinal nucleus of the trigeminal, oral part |
| STRv | Striatum ventral region |
| sAMY | Striatum-like amygdalar nuclei |
| SLC | Subceruleus nucleus |
| SPA | Subparafascicular area |
| SPF | Subparafascicular nucleus |
| SI | Substantia innominata |
| SCm | Superior colliculus, motor related |
| SCiw | Superior colliculus, motor related, intermediate white layer |
| Su3 | Supraoculomotor periaqueductal gray |
| SO | Supraoptic nucleus |
| SUT | Supratrigeminal nucleus |
| TEa | Temporal association areas |
